## Supplemental Figure 1 for "Priority effects alter interaction outcomes in a legume-rhizobium mutualism"

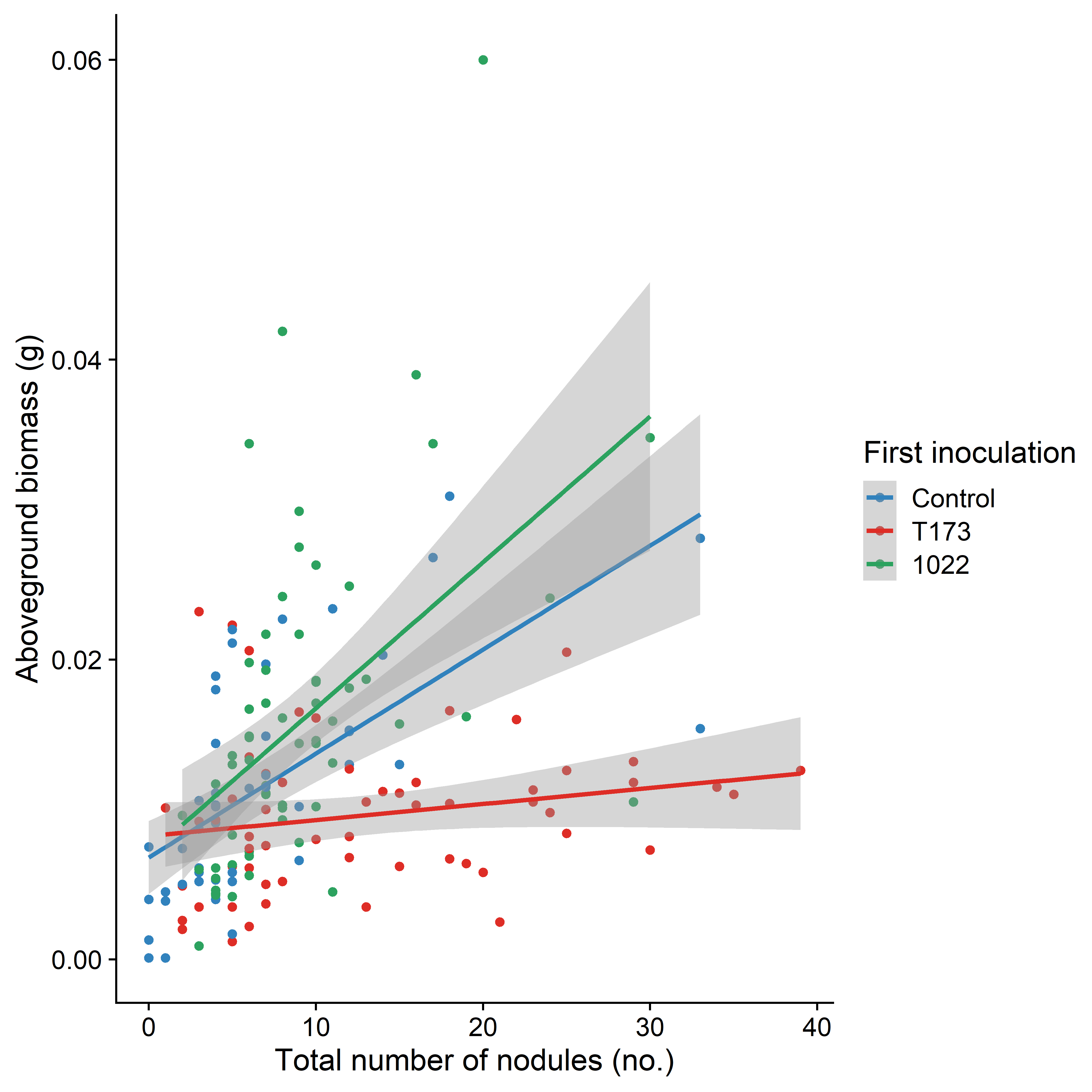

*Supplemental Figure 1. Effect of nodules on plant performance*: As nodulation increases, performance increases as well, but this increase is dampened when plants receive the ineffective strain first. Linear regression lines are by first inoculation, and include 95% confidence intervals in grey. Treatments are coded by order of strains, mutualistic *Ensifer meliloti* 1022, ineffective *Ensifer* sp. T173, and control inoculum. Colour indicates the first strain.
