## Supplemental Figure 2 for "Priority effects alter interaction outcomes in a legume-rhizobium mutualism"

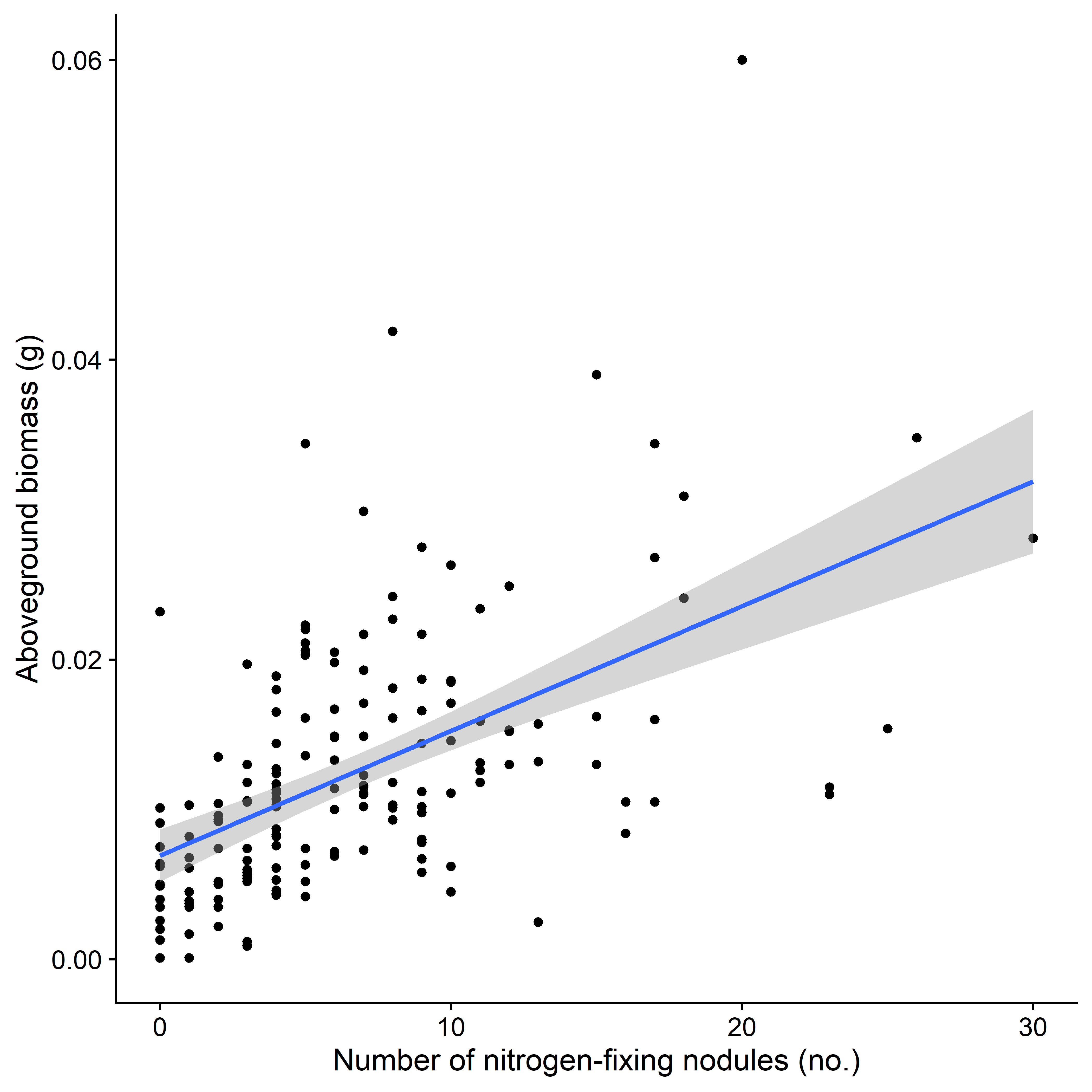

*Supplemental Figure 2. Correlation of Nitrogen-Fixing Nodules to Plant Performance:* There is a significant positive correlation between the number of nodules identified as nitrogen fixing and aboveground biomass (g).
