## Supplemental Figure 3 for "Priority effects alter interaction outcomes in a legume-rhizobium mutualism"

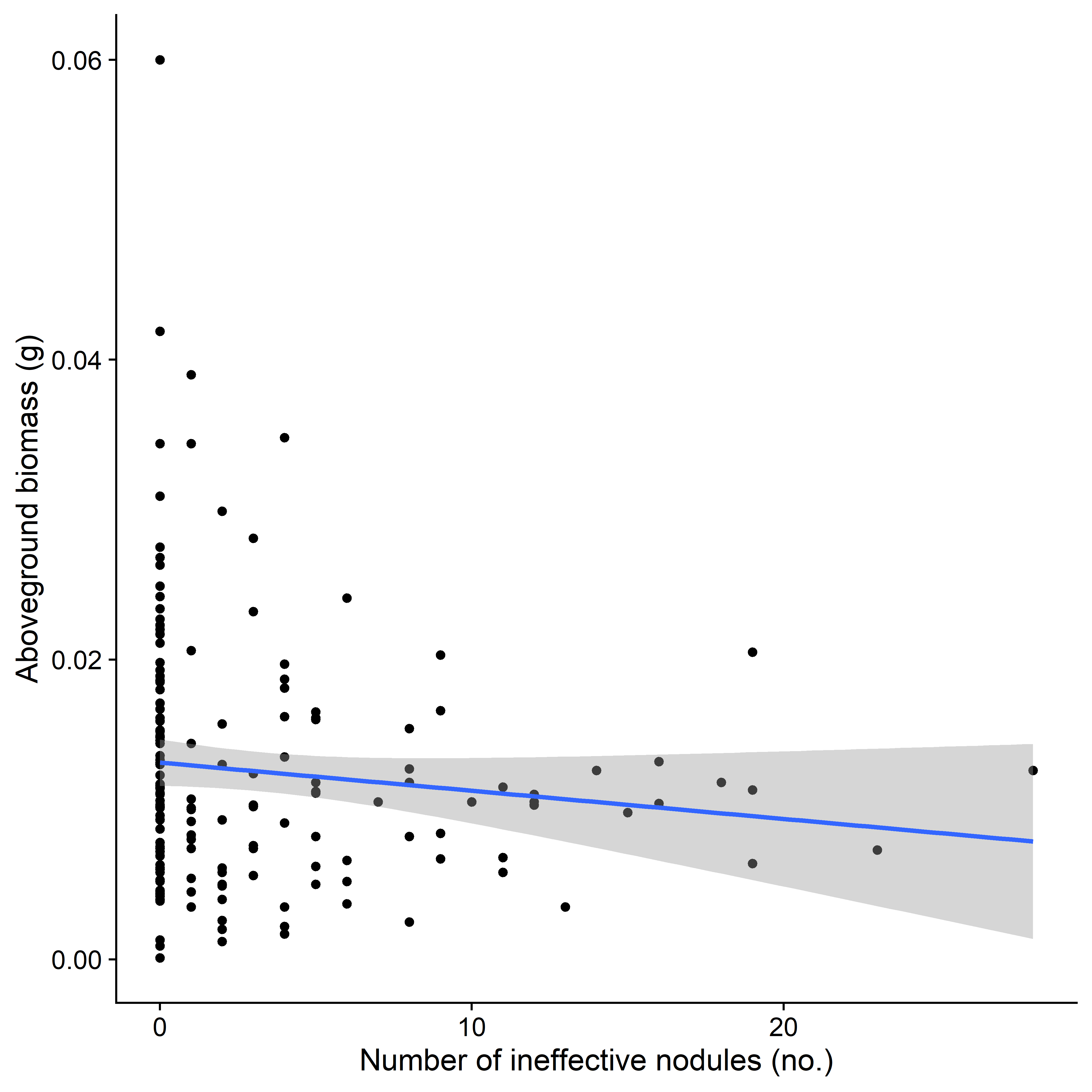

*Supplemental Figure 3. Correlation of Ineffective Nodules to Plant Performance:* There is a non-significant negative correlation between the number of nodules identified as ineffective and above ground biomass (g).
