## Supplemental Figure 4 for "Priority effects alter interaction outcomes in a legume-rhizobium mutualism"

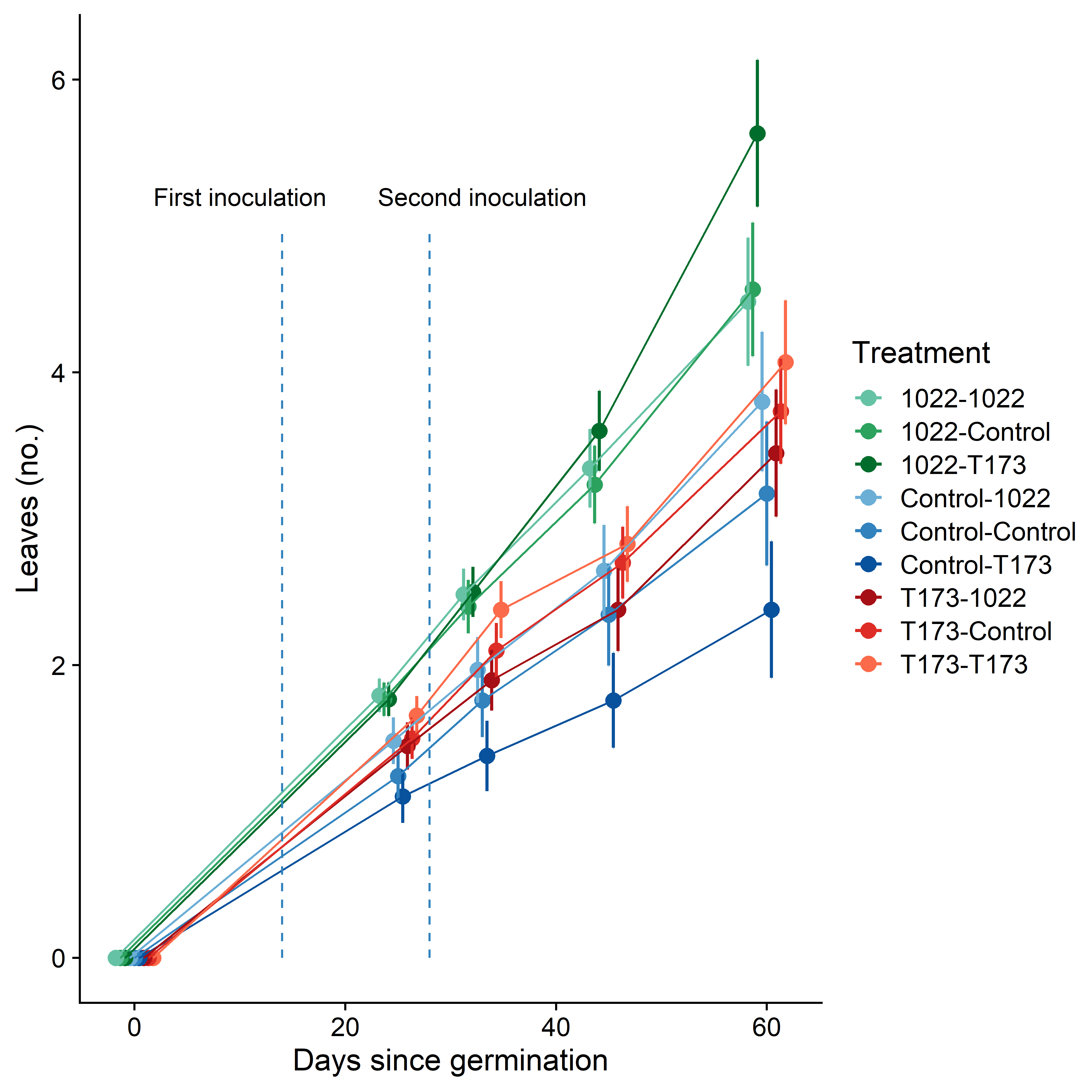

*Supplemental Figure 4. Effect of treatment on plant leaf number through time*: Shown here is the total number of leaves plants produced during the experiment through time. Note that after the first inoculation, rank order of treatment does not change much from the second inoculation. Treatments are coded by order of strains, mutualistic *Ensifer meliloti* 1022, ineffective *Ensifer* sp. T173, and sham inoculations (control). Colour indicates treatment and first inoculation.
