## Supplementary Material for "Priority effects alter interaction outcomes in a legume-rhizobium mutualism"

Fertilizer recipe: mix all ingredients in 500ml of distilled water, autoclave, wait for it to cool and then give each plant 1000ul.

MgSO_4_     2ml

NaCl          0.2ml

K_2_HPO_4_    1.7 ml

K2SO4       1.384 ml

CaCl_2_        2.4765 ml

Ca(NO_3_)    0.235 ml

KNO_3_        0.08 ml

FeEDTA    1.25 ml

MnSO_4_    0.050 ml

CuSO_4_     0.050  ml

ZnSO_4_     0.050 ml

H_3_BO_3_    0.050 ml

Na_2_MoO_4_  0.050 ml
